# Tagmentation-Based Mapping (TagMap) of Mobile DNA Genomic Insertion Sites

**DOI:** 10.1101/037762

**Authors:** David L. Stern

**Affiliations:** Janelia Research Campus, Howard Hughes Medical Institute, Ashburn, VA 20147, USA

## Abstract

Multiple methods have been introduced over the past 30 years to identify the genomic insertion sites of transposable elements and other DNA elements that integrate into genomes. However, each of these methods suffer from limitations that can frustrate attempts to map multiple insertions in a single genome and to map insertions in genomes of high complexity that contain extensive repetitive DNA. I introduce a new method for transposon mapping that is simple to perform, can accurately map multiple insertions per genome, and generates long sequence “reads” that facilitate mapping to complex genomes. The method, called TagMap, for <u>Tag</u>mentation-based <u>Map</u>ping, relies on a modified Tn5 tagmentation protocol with a single tagmentation adaptor followed by PCR using primers specific to the tranposable element and the adaptor sequence. Several minor modifications to normal tagmentation reagents and protocols allow easy and rapid preparation of TagMap libraries. Short read sequencing starting from the adaptor sequence generates oriented reads that flank and are oriented toward the transposable element insertion site. The convergent orientation of adjacent reads at the insertion site allows straightforward prediction of the precise insertion site(s). A Linux shell script is provided to identify insertion sites from fastq files.

## Introduction

Over the past approximately 30 years, multiple methods have been introduced to map the genomic insertion sites of transposable elements (Table 1). The original, and most widely used, technique, inverse PCR (iPCR), involves fragmentation of genomic DNA with restriction enzymes, followed by circularization of small DNA fragments and PCR with divergent primers specific to the transposable element to amplify DNA adjacent to the insertion site (Ochman *et al.* 1988). This method has been widely used, but requires use of restriction enzymes that both cut within the transposable element and ensure that the circularized molecules are small enough to allow PCR amplification. This sometimes leads to relatively short sequence reads that can be difficult to map unambiguously in complex genomes of higher eukaryotes. A second method, called splinkerette PCR, was developed to address this limitation (Devon *et al.* 1995) and has been used in a wide variety of applications to map transposable element insertion sites (Horn *et al.* 2007; Uren *et al.* 2009; Potter and Luo 2010). This method also involves fragmentation with restriction enzymes followed by ligation of a splinkerette oligonucleotide linker that contains a stable hairpin loop. PCR is employed to amplify DNA flanking the transposable element. Since splinkerette does not require restriction digestion of DNA within the transposable element, it is possible in principle to isolate longer sequences adjacent to inserted transposable elements.

**Table 1.** Comparison of methods for mapping transposable element genomic insertion sites.

| Method | Requires restriction digestion | Requires ligation step | Requires DNA shearing | Can map multiple insertions (without cloning) | Can map multiple non-unique insertions | Generates long reads for unambiguous mapping |
| --- | --- | --- | --- | --- | --- | --- |
| Inverse PCR | Yes | Yes | No | No | No | Sometimes |
| Splinkerette | Yes | Yes | No | No | No | Usually |
| Splinkerette-seq | Yes | Yes | No | Yes | Yes | Usually |
| Gohl *et al.* | No | Yes | Yes | Yes | No | Yes |
| TagMap | No | No | No | Yes | Yes | Yes |

Despite these improvements, both iPCR and splinkerette PCR still have several limitations. First, if a genome contains multiple transposable element insertions, then both methods may fail either because they amplify sequence adjacent to only one of the transposable elements or because both regions are amplified and Sanger sequencing generates uninterpretable results. It is also possible to perform high throughput sequencing on splinkerette products to circumvent this limitation (Wang *et al.* 2007; Uren *et al.* 2009), but this involves preparing a sequencing library in addition to the somewhat lengthy splinkerette protocol. Second, since both iPCR and splinkerette PCR depend on restriction digestion, it is impossible to guarantee recovery of sufficient sequence adjacent to all transposable element insertions.

Gohl *et al.* introduced a method that circumvents many of these limitations (Gohl *et al.* 2014). In this method, multiple pooled samples are generated and processed for next generation sequencing, so that each source strain is associated with a particular digital encoding (presence/absence) in each pool. The insertion site for each strain is then detected as a particular presence/absence pattern in the sequenced pools. This method appears to work well when each transposable element insertion site in the library is unique. The encoding would not be unique, however, if multiple strains carry elements inserted in the same location. I have found that piggyBac (pBac) transposable elements often transpose into genomes of non-model *Drosophila* species with high efficiency, leading to strains containing multiple insertion events. In the process of genetically parsing out these insertions, many of our strains contain the same insertion events. In this case, it is unlikely that insertion sites could be unambiguously mapped back to the original strains with the Gohl *et al*. method.

Given these considerations, it would be useful to have a method that can detect multiple transposable elements in a single strain and where the products of individual strains can be identified unambiguously. Here I describe a simple method that allows detection of all transposable elements in each of many strains. The method exploits tagmentation by Tn5 transposase, which simultaneously fragments DNA and attaches adaptors onto the ends of these fragments. This is analogous to performing restriction digestion and adaptor ligation in a single step, except that tagmentation generates DNA breaks at random positions. These random tagmentation positions offer the opportunity to generate extensive sequence data adjacent to insertion sites, which improves mapping accuracy. Unlike classical tagmentation, where two different adaptors are introduced simultaneously, TagMap uses only a single tagmentation adaptor that corresponds to the end that will be sequenced. As an example, I employ adaptor sequences for Illumina sequencing, but the adaptors for any high-throughput sequencing platform could be used. To isolate DNA adjacent to transposable elements, primers specific to both ends of the transposable element and one primer specific to the adaptor sequence are used during PCR of tagmentation products. During Tn5 transposition, only the 3’ end of the adaptor sequence is transferred to the target DNA (Whitfield *et al.* 2006). Thus, during normal library preparation methods, prior to thermocycling during PCR, the 3’ ends of tagmented fragments are first extended using a non-hot start DNA polymerase to generate the double strand adaptor sequence. In TagMap, we suppress creation of this double stranded adaptor sequence in all tagmented products in the reaction except those products that contain our target sequence. To accomplish this, we perform a hot-start PCR. In addition, we use a Tn5 adaptor in which the “top”, non-transferred, adaptor sequence contains an inverted thymine at the 3’ end, to suppress polymerization of primer dimers. Thus, the vast majority of DNA strands that contain a 3’ adaptor sequence suitable for PCR amplification will be products of first strand synthesis from the primers specific to the transposable element. I also show how barcodes can be included in the PCR primers, which can allow very high levels of multiplexing in a single library. Sequencing this library results in short reads that flank, and are oriented toward, the transposable element insertion site. TagMap produces sequencing reads up to approximately 800bp flanking the transposable element on each side, allowing unambiguous identification of genomic insertion sites in most cases.

## Methods and Materials

DNA was prepared from individual or pooled samples of adult *Drosophila* flies using the Quick-gDNA Micro Prep kit (Zymo). DNA was eluted to a concentration of approximately 20 ng / uL. Tn5 transposase was prepared according to the protocol described by Picelli *et al*. (Picelli *et al.* 2014). Tn5 transposase concentration was titrated to generate tagmentation of 20 ng *Drosophila* gDNA with adaptor A to an average size of approximately 500 bp. The size distribution of tagmented products was assayed by performing non-hot start PCR, with an initial extension step prior to denaturation, for 30 cycles with a primer complementary to the A adaptor and examining the distribution of reaction products on an agarose gel or an Agilent 2100 Bioanalyzer. This calibration procedure was performed only once per batch of Tn5 transposase.

Normal tagmentation protocols include a strand extension step to generate the complementary strands of the adaptors. However, for TagMap, we want to minimize creation of these complementary sequences from the original tagmentation products. Therefore, I used the high fidelity hot start thermostable polymerase Phusion polymerase (NEB) to favor first strand synthesis from the targeting primers.

TagMap uses divergent PCR primers complementary to the 5’ and 3’ ends of a targeted sequence and one primer complementary to the adaptors that are added during tagmentation. To block amplification of primer dimers from the non-attached top strand of tagementation adaptors, these oligonucleotides were synthesized with an inverted thymine at the 3’ end.

The protocol is illustrated in Figure 1, the sequences of all DNA adaptors and primers are provided in Supplementary Table 1, and a detailed protocol is provided as supplementary material. Tn5 protein was produced following the protocol provided in Picelli *et al.* (Picelli *et al.* 2014) and, just prior to tagmentation, was diluted to 20ng/uL in reassociation buffer (10 mM Tris pH 8.0, 50 mM NaCl, 1 mM EDTA). The Tn5 adaptor sequences Tn5ME-A and Tn5MErev were annealed in reassociation buffer by mixing 10uL of each oligonucleotide at an original concentration of 100uM in 80 uL of reassociation buffer. Oligonucleotides were annealed with the following thermocycler program: 95°C for 10 min, then 1 min at 90°C, and the temperature was decremented by 1°C for 60 cycles and the sample held for 1 min at each temperature. The annealed adaptors were diluted to 1 uM in H2O. The Tn5 was pre-charged with adaptors by thoroughly mixing 21 uL of 20 ng/uL Tn5, 10 uL glycerol, and 10 uL of 1 uM adaptors and incubating the mixture at 37°C for 30 min. Twenty ng of *Drosophila* gDNA was tagmented by mixing 1 uL of precharged Tn5, 1 uL of 20 ng/uL gDNA, 2 uL of 5 X TAPs buffer (50 mM TAPS-NaOH pH 8.5, 25 mM MgCl2, 50% v/v dimethylformamide, pH 8.5 at 25°C) and water to a total volume of 10 uL and then by incubating the mixture at 55°C for 7 min. 2.5 uL of 0.2% SDS was added to each reaction, which was then incubated at 55°C for 7 min to inactivate the Tn5 and release it from the DNA.

**Figure 1.**
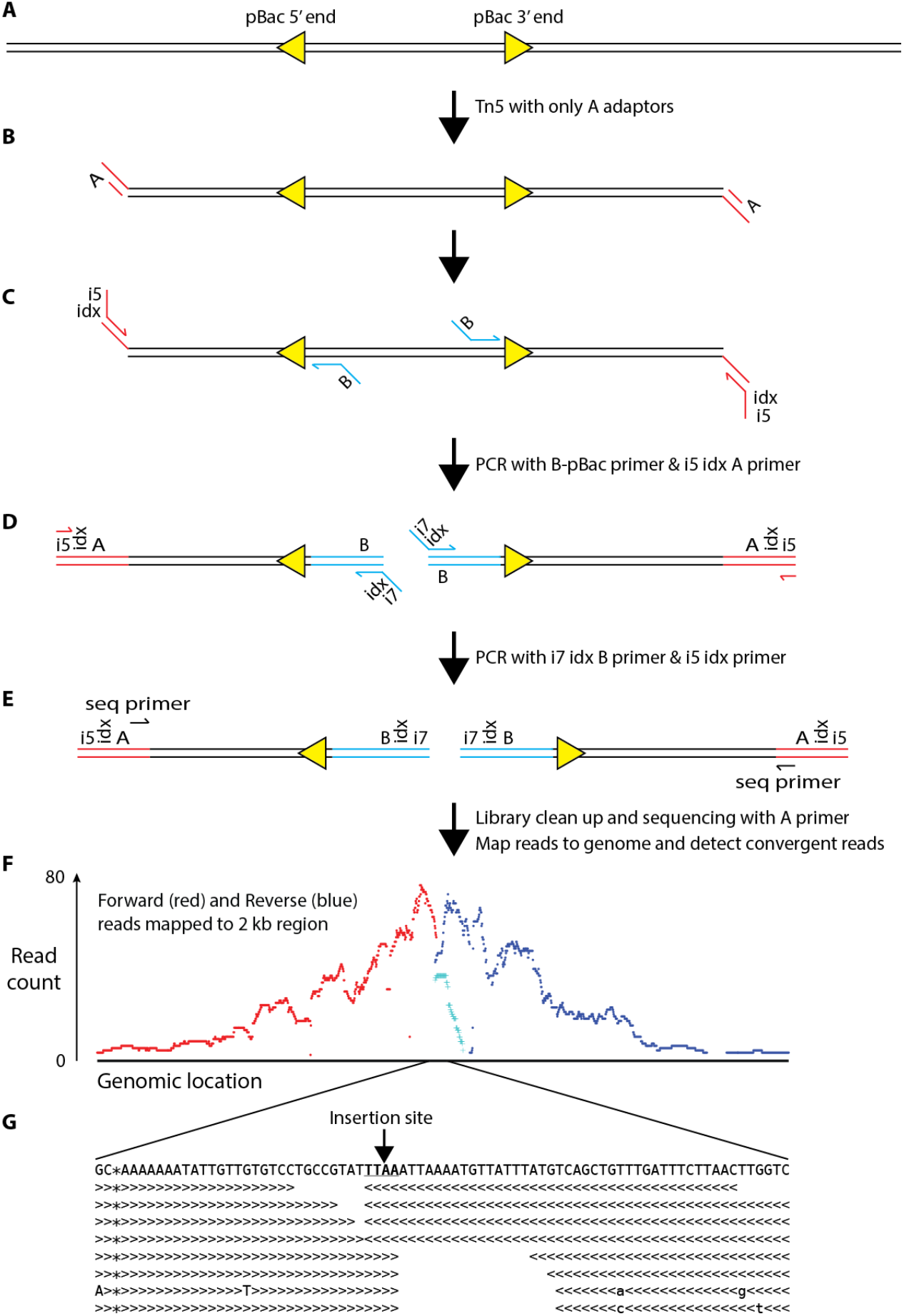
Schema of the TagMap protocol and representative results. (**A**) A genomic region containing a pBac transposable element is illustrated. DNA is tagmented with Tn5 protein pre-charged with a single A adaptor. (B) The tagmented products carry an A adaptor attached at their 3’ ends. (C) PCR with divergent primers internal to the transposable element and primers complementary to the A adaptor and carrying indexes (idx) and i5 sequences results in preferential amplification of sequences flanking the transposable element. (D) The i7 adaptors are added during a second PCR, which can also include a second set of indexes for higher levels of multiplexing. (E) The library is cleaned to remove residual primers, size selected, and sequenced with primers complementary to the A-i5 adaptor. The resulting sequencing reads are mapped to the genome sequence and insertion sites are detected as regions with adjacent, convergent reads. (F) Sequencing generates convergent reads flanking the insertion site. Coverage from forward and reverse reads for one of the pBac elements mapped in this study is shown in red and blue, respectively. Reads indicated in light blue overlap the 5’ end of the pBac plasmid, indicating the plasmid orientation (reverse in this case). (G) Examination of the region where forward (>) and reverse (<) reads overlap reveals the precise insertion site for this example at the canonical TTAA sequence for pBac elements. Only a small subset of the forward and reverse reads are shown.

For the first round of PCR, 1 uL of the stopped tagmentation reaction, 1 uL of 5uM targeting primer, 1 uL 5uM A_idx_i5 primer, 1 uL 4 mM dNTPs, 4 uL 5X Phusion Reaction Buffer, 0.5 uL Phusion Polymerase and 11.5 uL H2O were combined. The final reaction volume was 20 uL. The reaction was heated to 95°C for 5 min and then thermocycled 20 times at 95°C for 15 sec, 60°C for 15 sec, and 72°C for 1 min, followed by a final extension of 2 min at 72°C.

The i7 adaptor sequence was added during a second PCR. I combined 1 uL of the first PCR reaction, 1 uL of 5uM FC2 primer, 1 uL 5uM B_idx_i7 primer, 1 uL 4 mM dNTPs, 4 uL 5X Phusion Reaction Buffer, 0.5 uL Phusion Polymerase and 11.5 uL H2O. The final volume was 20 uL. The reaction was heated to 95°C for 5 min and then thermocycled 20 times at 95°C for 15 sec, 60°C for 15 sec, and 72°C for 1 min, followed by a final extension of 2 min at 72°C. Alternatively, for multiplexing, 10 uL from each of multiple samples from the first PCR can be pooled and 1 uL of this pooled sample used in the second PCR.

The products of PCR2 were cleaned and size selected using a 0.8:1 ratio of Ampure beads (Agencourt) to PCR reaction and then quantified on an Agilent 2100 Bioanalyzer. The library was sequenced on an Illumina HiSeq 2500 to generate 100 bp reads and the i5 indexes.

A Linux shell script was assembled that performs the following bioinformatics steps. (1) Duplicate PCR reads are filtered out using *prinseq* (Schmieder and Edwards 2011). (2) Unique reads are mapped to the genome sequence using *bwa-mem* (Li 2013). (3) Regions of the genome with convergent overlapping or adjacent forward and reverse reads are identified using *samtools* (Li *et al.* 2009) and various Linux commands. Most transposable elements duplicate sequences at the insertion site, such as pBac in this study, and insertion sites can therefore be detected as sites where forward and reverse reads overlap. For transposable elements that do not duplicate the insertion site, and for genome duplications, the script detects directly adjacent forward and reverse reads. (4) Only regions with more than one forward and reverse read are retained. (5) The orientation of the transposable element is determined by identifying reads that include both genomic and transposable element DNA. (6) Maps of the regions surviving these quality control steps are printed out at low and high resolution. Insertion sites in these images are easily identified because low-resolution images contain an obvious pileup of forward reads on one side and reverse reads on the other and the high-resolution view (using the *tview* command from samtools) allows identification of the precise insertion site (Figure 1f, g).

## Results and Discussion

The TagMap method and a representative result are shown in Figure 1. The method requires, first, a short tagmentation step using only a single Tn5 adaptor (Figure 1a,b). The use of a single adaptor differs from all other published tagmentation protocols and also means that commercial tagmentation kits cannot be used. Therefore, this method requires use of Tn5 free of adaptors. Fortunately, Tn5 can be produced using a straightforward protocol that was published recently by Picelli *et al.* (Picelli *et al.* 2014). In our hands, their published method works well and produces Tn5 with effectively no bacterial DNA contamination.

Tagmentation is followed by PCR with primers specific to the transposable element and oriented away from the element and toward the genomic DNA (blue primers in Figure 1c). The reverse primers are specific to the A adaptors that were added during tagmentation and contain sequences suitable for a high-throughput sequencing platform together with an optional index (barcode) to allow multiplexing of samples. A second index can be used in the i7 primer during a second PCR. Use of dual barcodes allows extremely high levels of multiplexing. Since relatively few sequencing reads are required to map a transposable element insertion site, it may be inefficient to use an entire lane of sequencing for this method. For mapping a small number of strains, I normally spike TagMap libraries into a more complex library at extremely low levels, usually at 0.01% or 0.1% of HiSeq libraries. It is easy to design barcodes that do not conflict with commercial kits, allowing identification of TagMap reads independently of reads from the main library. In Supplementary Tables 2 and 3, I provide a list of 96 i5 and i7 indexed primers for Illumina sequencing that do not conflict with the barcodes provided in Illumina kits.

Sequencing is performed only from the i5 adaptor sequence (Figure 1e). Informative reads are therefore oriented toward the insertion site (Figure 1f). The precise insertion site and transposable element orientation can be inferred from reads that overlap the insertion site (Figure 1f,g).

I tested TagMap by performing the protocol on multiple *Drosophila* strains carrying pBac elements. In every case, TagMap identified the correct previously identified insertion site for a pBac element. I tested whether TagMap can identify multiple elements in a single strain by recombining two previously mapped elements onto a single chromosome. TagMap correctly identified the locations of both pBac elements in this strain. In a separate case, previous iPCR experiments had indicated that one strain had only one pBac element inserted, but TagMap indicated that there were two pBac elements in this strain. I confirmed this result with specific PCR of both elements (data not shown). Therefore, TagMap can accurately map two transposable elements in a single strain where iPCR identified only one. The two elements have apparently remained associated in this single strain because they are located close to, and flank, the centromere of one chromosome.

To further explore the performance of TagMap, I examined the distribution of mapped reads around the insertion sites. First, I inserted the DNA sequence of the pBac plasmid into the genome sequence at the site identified by TagMap and remapped the TagMap reads (Figure 2). This revealed that sequence data is generated from the primers outward up to about 1 kb. This limit most likely reflects limitations of the Illumina sequencing platform, which normally cannot cluster amplicons longer than 1 kb. I then examined the distribution of reads for all nine mapped elements (Figure 3) and found that TagMap generates 1631 ± 167 bp (mean ± 1 SD) of data for each insert. This quantity of sequence should be sufficient to uniquely map mobile elements in the vast majority of cases. This experiment allowed me to estimate the fraction of all sequence reads that map to the insertion site. I found that 3.4% of reads mapped to this insertion site. That is, the majority of reads map to other regions of the genome. It is possible that the fraction of accurately mapped reads could be improved with optimization of the PCR steps, including reducing the number of PCR cycles and increasing the annealing temperature. In any case, this result emphasizes the importance of performing a bioinformatic analysis of the sequence data that searches for the identifying marks of a true transposon insertion site, including the presence of convergent sequencing reads that overlap the insertion site.

**Figure 2.**
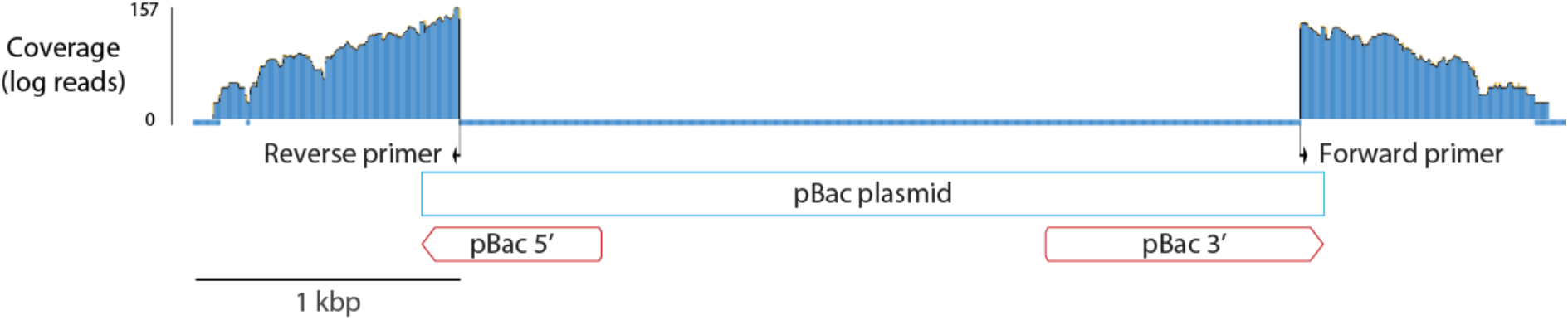
Coverage of sequencing reads mapped to a genomic region with a pBac plasmid inserted at the site identified by TagMap. Sequencing reads extend from the locations of the primers outward approximately 900 bp. The two pBac primers used for TagMap are represented as arrows and labeled.

While I have demonstrated mapping of artificially inserted transposable elements, it is easy to imagine several other applications of TagMap. First, naturally occurring transposable elements often display population variation for insertion sites and it is challenging to map these insertion sites. It is possible that TagMap can assist in the mapping of these mobile elements. Second, whole genome re-sequencing often reveals regions with an approximate doubling of sequence read depth, which may represent duplication events. It has been challenging to identify the genomic locations of these events. It is possible that TagMap can facilitate clarification of these genomic events.

**Figure 3.**
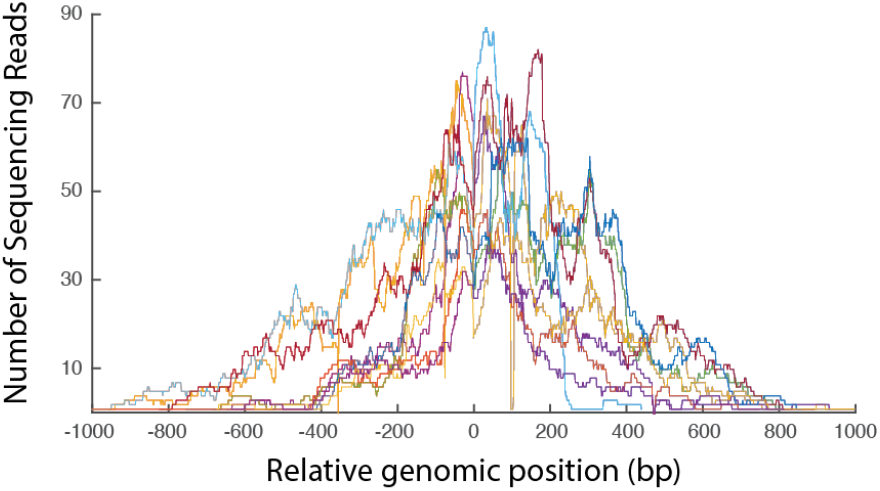
Density of read coverage flanking the insertion sites for nine transposable elements. The insertion site is position 0 and 1000 bp upstream and downstream of the insertion sites are shown. Separate insertion sites are shown in different colors. In all cases, reads to the left of 0 are forward reads and reads to the right are reverse reads. On average, the reads for each insertion site span 1631 ± 167 bp (mean ± SD).

## Acknowledgements

I am very grateful to Serge Picard for making the Tn5 protein used in these experiments and for sequencing the libraries. The development of TagMap benefited tremendously from discussions with Serge Picard and Andy Lemire over several years. I thank Yun Ding, Daryl Gohl and Andy Lemire for helpful comments on the manuscript. I am grateful to my lab members for ignoring me when I am puttering around in the lab.

